# Genome-wide development of intra- and inter-specific transferable SSR markers and construction of a dynamic web resource for yam molecular breeding: Y2MD

**DOI:** 10.1101/2023.03.09.531889

**Authors:** Moussa Diouf, Yedomon Ange Bovys Zoclanclounon, Pape Adama Mboup, Diaga Diouf, Erick Malédon, Ronan Rivallan, Hâna Chair, Komivi Dossa

**Affiliations:** Département de Mathématiques et Informatique, Faculté des Sciences et Techniques, Université Cheikh Anta Diop, BP 5005 Dakar-Fann, 10700, Dakar, Senegal.; Laboratoire Campus de Biotechnologies Végétales, Département de Biologie Végétale, Faculté des Sciences et Techniques, Université Cheikh Anta Diop, BP 5005 Dakar-Fann, 10700, Dakar, Senegal.; Department of Crop Science and Biotechnology, Jeonbuk National University, Jeonju 54896, Republic of Korea.; CIRAD, UMR AGAP Institut, 97170 Petit Bourg, Guadeloupe, France.; UMR AGAP Institut, Univ Montpellier, CIRAD, INRAE, Institut Agro, F-34398 Montpellier, France.; CIRAD, UMR AGAP Institut, F-34398 Montpellier, France.

**Author notes:** Correspondence: Komivi Dossa. These authors contributed equally to this work (Moussa Diouf, Yedomon Ange Bovys Zoclanclounon).

**Keywords:** genotyping, *Dioscoreacea*, database, microsatellite, transferable markers

## Abstract

**Background:** Microsatellite markers represent a low-cost and efficient tool for rapid genotyping as compared to single nucleotide polymorphism markers in laboratories with limited resources. For the economically important yam species widely cultivated in developing countries, very few microsatellite markers are available and no marker database has been developed to date. Herein, we conducted a genome-wide microsatellite marker development among four yam species, identified cross-species transferable markers, and designed an easy-to-use web portal for the yam breeder community.

**Results:** The screening of yam genomes resulted in 318,713; 322,501; 307,040 and 253,856 microsatellites in *Dioscorea alata*, *D. rotundata*, *D. dumetorum*, and *D. zingiberensis*, respectively. Mono-, di- and tri-nucleotides were the most important types of repeats in the different species and a total of 864,128 primer pairs were designed. Furthermore, we identified 1170 cross-species transferable microsatellite markers. Among them, a subset of 17 markers were experimentally validated with good discriminatory power regarding the species and the ploidy levels. Ultimately, we created and deployed a dynamic Yam Microsatellite Markers Database (Y2MD) available at http://yamdb.42web.io/. Y2MD is embedded with various useful tools such as JBrowse, Blast, *insilico*PCR, and SSR Finder to facilitate the exploitation of microsatellite markers in yams.

**Conclusions:** The present work is the first comprehensive microsatellite marker mining across several yam species and will contribute to advance yam genetic research and marker-assisted breeding. The released user-friendly database constitutes a valuable platform for yam breeders, especially those in developing countries.

## Introduction

Microsatellites, also termed simple sequence repeats (SSRs) are tandemly repeated DNA sequences generally spanning 1 to 6 bp (Ellegren, 2004). SSRs are ubiquitous in the genome of higher organisms and exhibit genetic polymorphism at inter- and intra-species levels (Bruford and Wayne 1993; Ellegren, 2004; Amiteye, 2021). Therefore, they have been widely used in population genetics (Chen et al., 2022; Sapkota et al., 2022;), genotyping, marker-assisted breeding, genome-wide association analysis (Li et al., 2014; Jha et al., 2021; Kim et al., 2021), forensic investigation (Li et al., 2021; Wang et al., 2022), evolutionary studies (Nowicki et al., 2021; Song et al., 2021; de Freitas et al., 2022), and conservation genetics (Rodríguez-Peña et al., 2018; Dong et al., 2022; Ming et al., 2022).

Considering the low level of difficulty regarding the technical aspect of setting an SSR-based genotyping core facility, the relatively inexpensive cost of reagents, and the level of throughput, SSRs markers are quite more affordable and implementable in a standard laboratory (Herrera and Ghislain, 2013), especially those from low-income countries. Besides, SSRs can exhibit good discriminatory power, are randomly present in the genome, and can be located in the genic region, enabling the genetic dissection of traits of interest (Ellegren, 2004). Microsatellites also exhibit co-dominance with Mendelian inheritance, facilitating their traceability within different breeding populations (Normann et al., 2018). Therefore, owing to their simple identification and high reproducibility, breeders have preferentially relied on SSRs because they are easier to use than single nucleotide polymorphism (SNP) markers, which demand expensive charges and facilities (Wang and Wang, 2016).

In plant breeding, the development of molecular marker databases has been extensively promoted in economically important plants such as *Sesamum indicum* (Dossa et al., 2017), *Vicia faba* (Mokhtar et al., 2020), *Nicotiana spp*. (Wang et al., 2018), legumes species (n = 13) (Duan and Kaundal, 2021), horticultural species (n = 112) (Song et al., 2021), and *Anemone coronaria* (Martina et al., 2022). Since the usage is based on a graphical user interface, the database offers the breeders, a user-friendly path for rapid identification of molecular markers without a pre-requisite expertise in command line scripting. A web database concentrates all information in one platform and assists the breeders to design markers related to a trait of interest.

Yams (*Dioscorea* spp.) represent an important staple food crop, feeding over 300 million people in tropical and subtropical regions (Mignouna et al., 2008). The biggest producer country is Nigeria with 70% of the global production, followed by Ghana, and Côte d’Ivoire (https://www.fao.org/faostat/en/FAOSTAT). The genus contains approximately 650 species (Virouel et al., 2018). As a cash crop, yams contribute to the income of more than 60 million people (Asiedu et al., 2010), with a production of 74 million tons and an economic market value of USD 21 billion estimated in 2020 (United Nations Statistics Division 2022). Besides, in West Africa where yam is the most cultivated in the so-called “yam belt” (from Western Cameroon to central Côte d’Ivoire) (Sarcelli et al., 2019), more than five million people directly benefit from yam culture (Mignouna et al., 2020); making this crop a valuable resource for food security and poverty alleviation in this part of the African continent (Darkwa et al., 2020).

To accelerate yam breeding programs, attention has been paid to the development of polymorphic DNA markers. Restriction Fragment Length Polymorphisms (RFLPs) were first developed to investigate the origin and phylogeny of cultivated Guinea yams and wild relatives by using chloroplast DNA (cpDNA) and nuclear ribosomal (rDNA) (Terauchi et al., 1992). Later on, Amplified Fragment Length Polymorphism (AFLP) and Random Amplified Fragment DNA (RAPD) were employed for genetic diversity assessment (Malapa et al., 2003, Egesi et al., 2007, Zannou et al., 2009), yam mosaic virus resistance markers identification (Mignouna et al., 2002a), and genetic linkage map construction (Mignouna et al., 2002b).

Furthermore, due to their high level of polymorphism and cost-effectiveness, SSRs were developed from cultivated and wild yam species (Terauchi and Konuma, 1994, Mignouma et al., 2003, Mizuki et al., 2005, Hochu et al., 2006, Tostain et al., 2006, Siqueira et al., 2011, Silva et al., 2014, Tamiru et al., 2015). However, certain yam species including *D. batatas* and *D. opposita* have not yet available SSR markers. More importantly, less than 100 validated SSR markers are available in yams to date and no information on their genome coverage is available. Knowing that yam breeding programs are mostly based in countries of the global South where SSRs are widely used (Tostain et al., 2007, Otto et al., 2015, Zawedee et al., 2014, Loko et al., 2016, Korsa et al., 2022, Nwogha et al., 2022), generating SSR markers will surely empower the breeders in their routine tasks such as paternity testing, hybrid identification, QTL detection, marker-assisted selection, genome-wide association study, etc.

Meanwhile, with the help of short-, long-reads and chromosome conformation sequencing technologies, good-quality genomes were recently released in yams. These data enable the identification of genomic regions associated with agronomically important traits such as sex determination systems (Tamiru et al., 2017, Cormier et al., 2019, Sugihara et al., 2020), anthracnose resistance, tuber oxidative browning, tuber starch content (Bredeson et al., 2022), post-harvest hardening (Siadjeu et al., 2021).

To date, the genomes of four species *v.i.z D. alata* (Bredeson et al., 2022), *D. rotundata* (Tamiru et al., 2017), *D. dumetorum* (Siadjeu et al., 2020), and *D. zingiberensis* (Cheng et al., 2021) are publicly available. The genomes presented a high level of collinearity (Bredeson et al., 2022), indicating that SSRs might be highly transferable from one species to another, even likely, to those wild species that have not yet been sequenced. Therefore, having a comprehensive SSR database for the yam breeders is fundamental for pre-breeding and breeding activities.

The present study was designed to (i) mine the genome of four yam species for the identification single-species and cross-species transferable SSR markers, (ii) create primer sets for the markers, (iii) evaluate the discriminatory power of selected markers, and (iv) build a digital web resource to support yam breeding and genetic research.

## Materials and Methods

### Genomic sequence resources

A total of four genome sequences were retrieved from the National Center for Biotechnology Information (NCBI) genome portal, including *Dioscorea alata* cultivar TDa95/00328 (GenBank assembly accession: GCA_020875875.1), *D. dumetorum* (GenBank assembly accession: GCA_902712375.1), *D. rotundata* cultivar TDr96_F1 (GenBank assembly accession: GCA_009730915.2), and *D. zingiberensis* isolate CJ2019 (GenBank assembly accession: GCA_014060945.1) (**Supplementary Table 1**). Besides, we added a newly assembled chromosome-scale assembly (unpublished data) of another greater yam (*D. alata*) “Kabusa”.

### Identification of microsatellites and primer design

SSRs were searched using the Genome-wide Microsatellites Analyzing Tool Package (GMATA) (Wang and Wang, 2016) and primers were designed from the flanking sequences (200 bp on each side) to probe the genomic sequences. The genome assembly was imported into the GMATA ‘SSR identification’ module in FASTA format. The microsatellite search parameters applied for SSR detection are: minimum length (nt) : 1, maximum length (nt) : 6, minimum number of repeats : 5.

Using custom python scripts, only sequences with a size greater than 10 bp were retained. The SSR locus information files generated (.ssr) contain the start and end positions of the SSRs on the chromosomal sequence, the SSR template and the number of repeat units.

The genome assemblies in FASTA format and the output files (.ssr) generated by the “SSR identification” module were imported into the “Marker Designing” module of GMATA. Primer design parameters were as follows: minimum amplicon size: 100 bp, maximum amplicon size: 400 bp, optimal annealing temperature: 60 °C, flanking sequence length: 200 bp, maximum model length (flanking sequence+SSR): 2000 bp. The output file (.seq) contains the repeats centred on 200 adjacent sequence bases on both sides. Output files with extensions .mk and .sts were created, containing left and right primer sequences, hybridization temperatures, primer positions on chromosomes and expected amplicon sizes.

The identified SSRs were classified into genic and non-genic SSRs for *D. alata* and *D. rotuntada* for which genome annotations are available. Subsequently, functional enrichment of genes containing SSRs was performed with KOBAS-i webserver (Bu et al., 2021) in order to find out their biological significance.

### *In silico* Polymerase Chain Reaction

The e-Mapping module was used to run the e-PCR algorithm (Schuler, 1997), to validate markers, generate amplicons and assign marker positions across the *Dioscorea* genomes. This module was also used for intraspecific (the two *D. alata* genomes) and interspecific marker mapping at the genome scale. In e-PCR, FASTA files containing all genome assemblies and newly developed markers created with the “Marker designing” module (.sts files) were imported as “Sequence File” and “Marker File ’’, respectively. The gap (-n) and max. indel (-g) parameters were set to 2 and 1, which means that two mismatches and one insertion are allowed. The output file (.emap) provides detailed marker amplification information with calculated amplification sizes and chromosomal targets and identifies single locus and multilocus markers, ultimately producing those that are potentially polymorphic (different sizes of PCR products).

## SSR primer validation

### Plant materials, DNA isolation and PCR experiment

Leaf samples from 10 yam species (*D. alata*, *D. rotundata*, *D. dumetorum*, *D. bulbifera*, *D. trifida*, *D. transversa*, *D. esculenta*, *D. cayenensis*, *D. abyssinica* and *D. nummularia*) including cultivated and wild species were used for DNA extraction. Six *D. alata* genotypes with different ploidy levels (14M, 74F, Pyramide, Kabusa (2x), CT148 and CT198 (4x)) and two D*. rotundata* (Dr1 and Dr2) were included (**Supplementary Table 2**). Plant samples are available in the germplasm collection of CIRAD, Guadeloupe. DNA was extracted based on the Mixed Alkyl Trimethyl Ammonium Bromide method (Cummings and Wood, 1989). Prior to the PCR experiment, the quantity and quality of the extracted DNA were consecutively assessed on 1% agarose gel and Invitrogen Qubit Flex Fluorometer. A total of 18 SSR primers from 18 chromosomes (**Supplementary Table 3**) which exhibited interspecific amplifications through the e-PCR were selected for the PCR experiment. M13 tail (CACGACGTTGTAAAACGAC) was added to the primers. The PCR reaction was performed using the QIAGEN kit under the conditions as followed 95 °C for 5 min, 10 cycles of 30 s at 95 °C, 60 °C for 1 min 30 s and 72 °C for 30 s, 25 cycles of 30 s at 95 °C for 1 min 30 s and 72 °C for 30 s, followed by 30 min at 60 °C. Migration of the PCR products was conducted on the ABI 3500xL (Thermo Fisher Scientific Inc.). Analysis of microsatellite profiles was performed with Genemapper v6.0 (Applied Biosystems™).

### SSR genotyping data analysis

The genotyping data served to perform an hierarchical clustering to depict the genetic relationships between the 16 accessions (**Supplementary Table 2**). The clustering was conducted in R v4.2.2 (R Core Team, 2022) following the Unweighted Pair Group Method with Arithmetic mean method using the function *hclust()* of the package base. The resulting tree was rendered with the function *fviz_dend()* of the package factoextra v1.0.7 (Kassambara and Mundt, 2020).

### Construction and deployment of Yam Microsatellite Markers Database

The Yam Microsatellite Markers Database (Y2MD) database conception relied on three major steps detailed in **Figure 1**: (i) data compilation in SQL database, (ii) back and front-end web construction, and (iii) tools integration.

**Figure 1.**
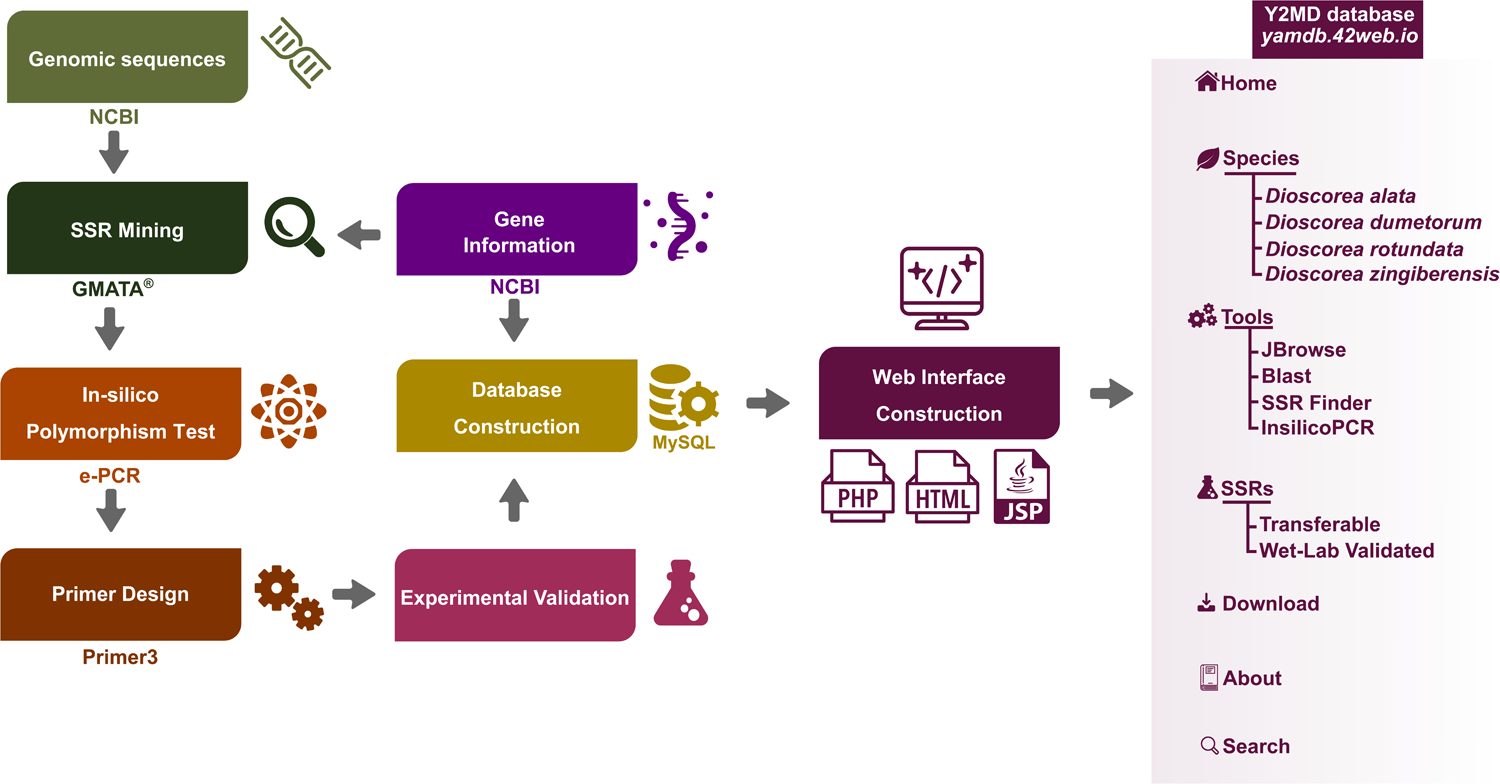
A diagram depicting the structure of Yam Microsatellite Markers Database (Y2MD).

In brief, using the generated SSR markers datasets, the Y2MD SQL database was built. Thus, the Y2MD server was implemented using Linux, Apache, MySQL, and PHP (LAMP) web application platform. The user-friendly web interface was then developed with the aid of JavaScript, and the HyperText Preprocessor (PHP). To enable the users to conduct their own analyses conveniently, additional functionalities such as JBrowse (Skinner et al., 2009), Blast (Altschul et al., 1990), SSR Finder (http://www.biophp.org/minitools/microsatellite_repeats_finder/, accessed on 15 March 2022), and InsilicoPCR (http://www.biophp.org/minitools/pcr_amplification/, accessed on 15 March 2022) were embedded. The website is freely accessible at http://yamdb.42web.io/.

## Results

### Detection, distribution and characteristics of microsatellites detected on yam genomes

The genome sequences of four yam species were used for the identification of simple sequence repeats (SSRs). In total, 318,713; 322,501; 307,040 and 253,856 SSRs were detected in *Dioscorea alata*, *D. rotundata*, *D. dumetorum* and *D. zingiberensis* genomes, respectively, with respective densities of 662.86 SSR/Mbp, 552.22 SSR/Mbp, 633.07 SSR/Mbp and 529.97 SSR/Mbp (**Figure 2; Supplementary Figure 1**). At the intraspecific level, the genome of the *D. alata* cultivar “Kabusa’’ contains 346,986 SSRs with a density of 693.972 SSR/Mbp, which is more than *D. alata* cultivar TDa95/00328.

**Figure 2.**
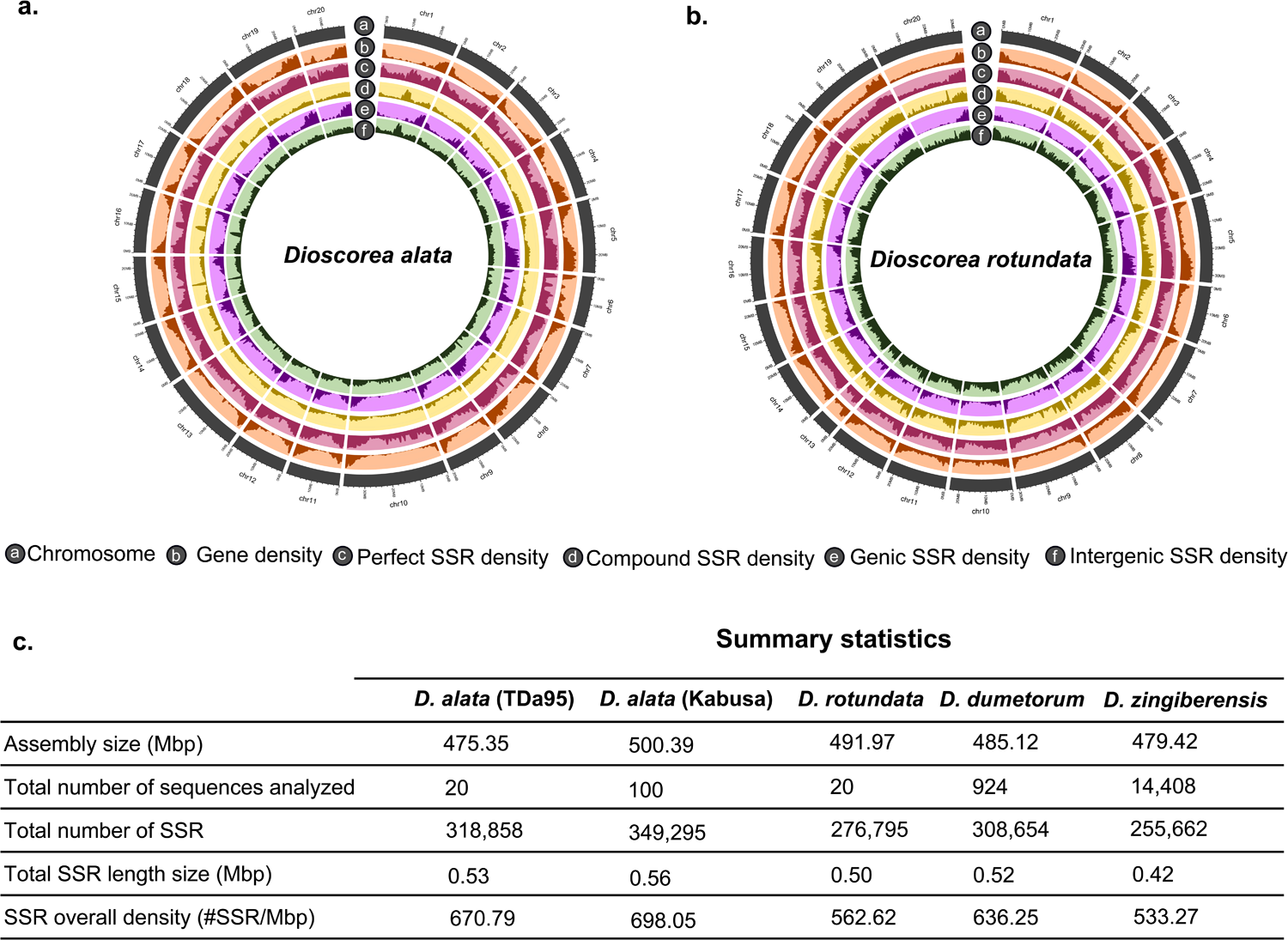
Distribution and summary statistics of SSRs identified in yam genomes. Circos plots depicting the SSR density within the chromosome-scale genomes of *Dioscorea alata* (TDa95) (a) and *Dioscorea cayenensis subsp. rotundata* (b). Summary statistics of SSR counts in the five studied genomes (c).

The numbers of SSRs per chromosomes are strongly correlated to the chromosome lengths (**Figure 3a**). A total of 658 different motifs were observed in *D. alata*, 843 in *D. dumetorum*, 762 in *D. rotundata* and 753 in *D. zingiberensis* (**Figure 3b**). Mono-, di- and tri-nucleotides are the most important types of repeats in the different species (**Figure 3c**). The motifs obtained are strongly rich in A/T: A and T mononucleotides are the most represented with an average of 55% of the detected SSRs, 20% of the SSRs are constituted by AT and TA di-nucleotides. For tri-nucleotides, repeats (AAT/ATA/ATT/TAA/TTA) account for 7.5%. Hexa- and penta-nucleotides had relatively low frequencies.

**Figure 3.**
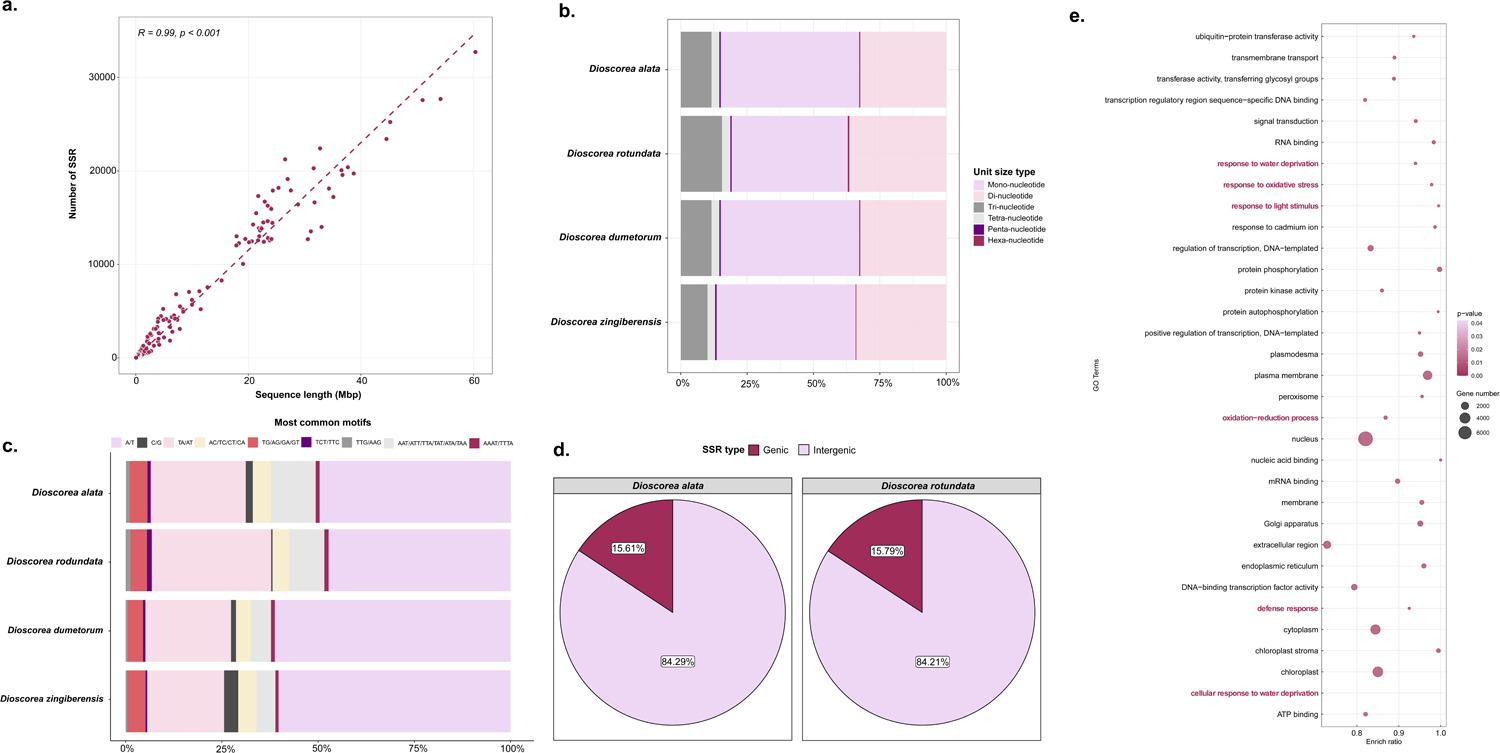
Correlation between chromosome length and number of SSRs (a); Proportions of the nucleotide repeats type (b); Distribution of the most common patterns in the four species (c); Proportions of genic and non-genic SSRs in *D. alata* and *D. rotundata* (d); Bubble plot showing significantly enriched gene ontology terms related to genic SSRs regions. The gene ontology terms highlighted in red are those that are agronomically relevant (e).

Most of the SSRs (∼84%) were localised in the intergenic regions in *D. alata* and *D. rotundata* genomes (**Figure 3d**). This analysis was not performed for other species because of the lack of a proper genome annotation. The genes containing SSRs were analysed for their functional enrichment. They were enriched in a wide panel of biological attributes. Among them, various agronomically relevant gene ontology terms were found including response to water deprivation (GO:0009414), response to oxidative stress (GO:0006979), response to light stimulus (GO:0009416), defense response (GO:0006952), and inflorescence development (GO:0010229) (**Figure 3e; Supplementary Table 4**).

### Development of genome-wide SSR primers

A total of 864,128 unique primer pairs were created from the 1,201,570 SSRs extracted from the four *Dioscorea* genomes: *D. alata* (221,606), *D. rotundata* (233,593), *D. dumetorum* (208,306) and *D. zingiberensis* (200,623). All primers are distributed unevenly across the genomes. The identified primers and their characteristics are presented in **Supplementary Table 5**. All the markers were assigned the identifiers YMa (*D. alata*), YMr (*D. rotundata*), YMd (*D. dumetorum*), YMz (*D. zingiberensis*) followed by the consecutive numbers of the primers in the genomes.

### Cross-species transferability of SSRs

First, we performed a pairwise comparison of the microsatellite markers between the four yam species (**Supplementary Table 6)**. *Dioscorea alata* and *D. rotundata* displayed the highest conservation rate (54,78%) while *D. rotundata and D. zingiberensis* had the lowest conservation rate (1,64%).

Next, to evaluate the cross-species transferability of the microsatellite markers, we set *D. alata* as a reference and performed an e-PCR on the other three species. Of the 221,606 SSRs markers from *D. alata* (TDa95/00328), 166,817 SSRs were specific to *D. alata*, while 39,837 SSRs were able to amplify *D. rotundata*, 3473 on *D. dumetorum* and 677 on *D. zingiberensis* (**Figure 4a**). Collectively we identified 1170 core SSR markers able to amplify all the four species and potentially more yam species.

**Figure 4.**
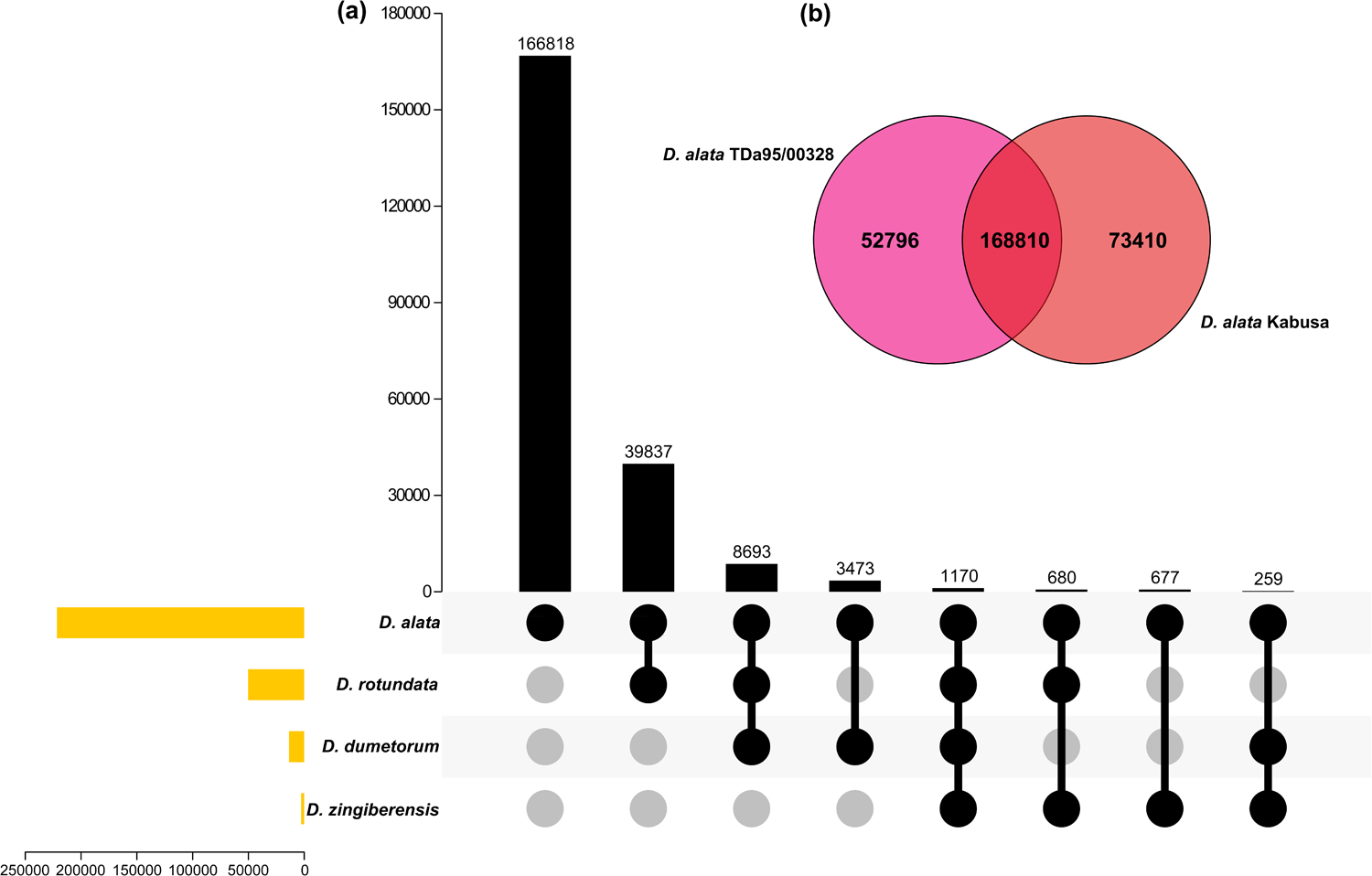
Cross-species and transferability of SSRs in yams. Upset plot showing the transferable markers between four yam species (*D. alata* as the reference genome, *D. rotundata*, *D. dumetorum* and *D. zingiberensis*) (a); Venn diagram showing the number of common and specific SSR markers detected in two *D. alata* cultivars (TDa95/00328 and Kabusa) (b).

At the intraspecific level, we compared the SSR maker data identified in the genomes of the two *D. alata* cultivars (TDa95/00328 and Kabusa). The results showed that both cultivars had in common 77% of the total SSR markers (**Figure 4b**).

### Validation of selected SSRs and genotyping analysis

Firstly, to validate the newly developed markers in yam species, we extracted 92 SSR markers from the literature routinely used for genotyping *D. alata* and *D. rotundata*. This validation was done using the e-PCR method. Of the 92 markers, 84 (91%) yielded PCR products on their respective reference genomes.

Next, a set of 18 interspecific markers were selected from 18 chromosomes and used for a wet-lab PCR validation in ten yam species (**Supplementary Table 2; Supplementary Table 3**). Out of the 18 selected markers, 17 produced amplicons (except the marker YMar128). Sixteen markers produced amplicons in at least six species over the ten (**Table 1**). Interestingly, the marker amplifications in diploid and tetraploid *D. alata* cultivars were in the range of maximum two or four alleles, respectively, suggesting the ability of those markers to potentially discriminate cultivars following the ploidy criteria. For instance the marker YMar79 was able to clearly delineate diploid from tetraploid forms of *D. alata* by the expression of the number of allele inferior or equal to the number of ploidy (**Table 1**). Interestingly, four alleles were amplified for *D. transversa* (markers YMar44, YMar79, and YMac82), *D. cayenensis* (markers YMar39, YMar44), *D. abyssinica* (markers YMar28 and YMac82), and *D. nummularia* (marker YMar79), indicating that the cultivars of these species used in this study are polyploids (4x or more). Besides, a maximum of two amplified alleles were found for both cultivars of *D. rotundata*, suggesting that they are diploid types.

**Table 1.** Allele scoring for 18 selected SSR markers in ten yam species

|  | *D. alata*<br>(14M) | *D. alata*<br>(74F) | *D. alata*<br>(CT148) | *D. alata*<br>(CT198) | *D. alata*<br>(Pyramide) | *D. alata*<br>(Kabusa) | *D. bulbifera* | *D. transversa* | *D. rotoundata* | *D. rotoundata* | *D. trifida* | *D. dumetorum* | *D. esculenta* | *D. cayenensis* | *D. abyssinica* | *D. nummularia* |
| --- | --- | --- | --- | --- | --- | --- | --- | --- | --- | --- | --- | --- | --- | --- | --- | --- |
| YMar02 | 198 | 199 | 197 + 199 | 197 + 199 | 198 + 199 | 196 + 197 | 212 + 213 | 198 + 199 | 198 + 199 | 198 | 215 | 209 | 204 | 198 + 199 | 197 + 200 | 196 + 199<br>+204 |
| YMar04 | 313 + 325 | 325 | 309 + 313 | 309 + 313 | 313 + 325 | 313 + 327 | 333 + 335 | 313 + 315 | 322 | 322 | 343 | 349 | 322 | 320 + 322 | 322 | 311 |
| YMar11 | 303 | 303 | 303 | 303 | 303 | 303 | 304 | 303 | 317 | 311 + 317 | 296 | 304 | 308 + 311<br>+ 317 | 303 | 314 | 302 |
| YMar118 | 196 | 202 | 202 | 202 | 196 | 202 | 194 | 202 | 198 | 196 | 214 |  |  | 197 | 197 | 198 |
| YMar128 | NA | NA | NA | NA | NA | NA | NA | NA | NA | NA | NA | NA | NA | NA | NA | NA |
| YMar135 | 160 | 160 | 157 + 160<br>+ 176 | 157 + 160 +<br>176 | 160 | 176 | 163 + 169 | 163 + 166<br>+ 172 | 160 | 160 | 176 | 160 + 163 | NA | 157 + 160 | 163 | 163 + 166<br>+ 172 |
| YMar15 | 158 | 158 | 158 + 159 | 158 + 159 | 158 | 158 | NA | 155 + 158 | 157 | 157 | 155 | 152 | 155 | 153 + 157 | 156 + 157 | 155 + 158 |
| YMar28 | 234 + 254 | 232 + 248 | 251 + 260 | 251 + 260 | 226 + 230 | 224 | 234 | 224 | 234 + 248 | 230 + 232 | 224 | NA | NA | 236 + 260 | 232 + 234<br>+ 238 +<br>240 | 224 |
| YMar33 | 181 + 183 | 183 + 184 | 181 + 184 | 181 + 184 | 183 | 184 | 170 + 189 | 176 + 184 | 184 | 184 | NA | 186 | 184 + 185 | 184 | 182 | 182 |
| YMar39 | 180 | 180 | 180 + 181 | 181 | 180 | 180 + 182 | 170 | 154 + 157<br>+ 181 | 173 + 176 | 176 | 166 | 171 | 185 | 173 + 176<br>+ 1778 +<br>186 | 173 | 154 + 157 |
| YMar44 | 192 + 206 | NA | 208 + 212 | 196 + 208 | 192 + 204 | 196 + 206 | NA | 184 + 192<br>+ 196 +<br>200 | 190 + 196 | 190 + 196 | NA | NA | NA | 190 + 194<br>+ 198 +<br>208 | 192 | 192 + 196 |
| YMar45 | 277 + 281 | 277 | 281 | 281 | 277 + 281 | 279 + 281 | NA | 275 + 277<br>+ 281 | 273 | 273 | NA | 281 | 259 | 267 + 275 | 275 | 275 |
| YMar59 | 298 | 298 | NA | NA | 298 | 298 | 310 | 298 + 301 | 307 + 310 | 310 + 319 | NA | NA | NA | 310 | 307 | 301 |
| YMar65 | 274 + 275 | NA | 274 + 277 | 274 + 277 | 275 | 273 + 275 | NA | 274 | NA | NA | NA | NA | NA | NA | NA | NA |
| YMar79 | 217 | 208 + 217 | 207 +<br>211+ 223 | 207 + 211+<br>220 + 223 | 217 + 223 | 220 | 198 | 207 +<br>211+ 221<br>+ 225 | 215 + 220 | 214 | 223 | 236 + 242 | 223 | 208 + 214 | 212 + 215 | 221 + 225+<br>229 + 236 |
| YMar82 | 238 + 254 | 238 + 254 | 226 + 238<br>+ 256 | 226 + 256 | 252 + 254 | 228 + 234 | NA | 230 + 238<br>+ 246 +<br>252 | 238 | 238 | NA | NA | NA | 238 + 248<br>+ 254 | 232 + 242<br>+ 246 +<br>264 | 236 + 246<br>+ 252 |
| YMar90 | 294 | 292 | 292 | 292 | 292 + 304 | 292 + 294 | 332 | 294 + 310 | 306 | 296 + 298 | 332 + 340 | NA | NA | 296 + 314 | 310 | 294 + 310 |
| YMar92 | 215 | 215 | NA | NA | 214 | 215 + 219 | 209 | 211 + 214 | 212 + 215 | 211 + 212 | NA | 215 + 223 | 219 | 211 + 212 | 216 | 211 |

Based on the genotyping data, we performed a hierarchical clustering analysis of the accessions and generated a tree. The tree highlighted a branch harbouring exclusively *D. alata* cultivars (**Figure 5**). Among *D. alata* cultivars, a clear distinction of diploid and tetraploid types was also noted, implying the robustness of the selected markers for discriminating yam accessions based on both species and ploidy criteria. The species *D. cayenensis* and *D. abyssinica* formed a distinct group individually while *D. transversa* and *D. nummularia* gathered together. Interestingly, the two cultivars of *D. rotundata* grouped together as expected. Meanwhile, two other groups were also pointed out: one consisting of *D. trifida* and *D. esculenta*, and another group formed by *D. dumetorum* and *D. bulbifera*.

**Figure 5.**
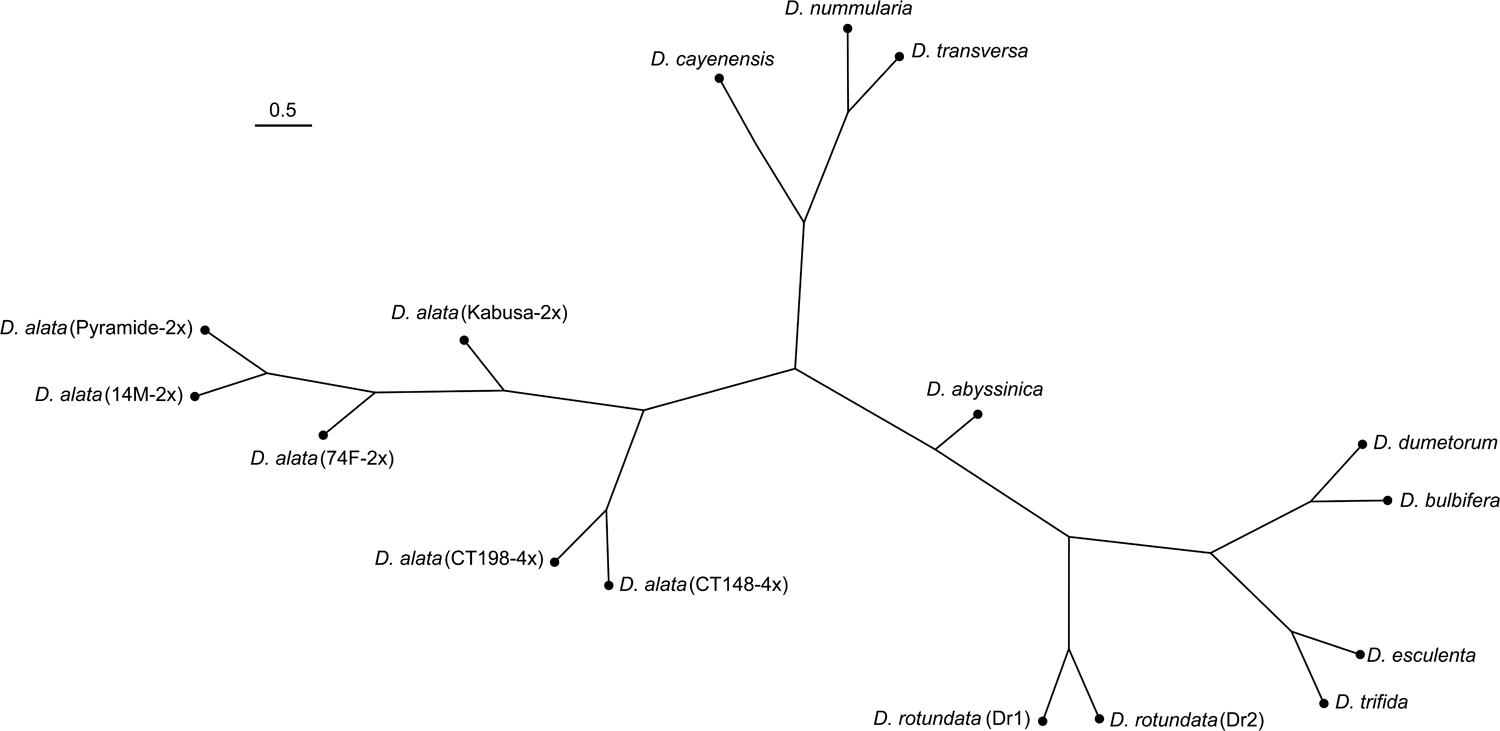
Hierarchical clustering tree depicting the relationships between ten yam species using 17 SSR markers. The tree was constructed based on the Unweighted Pair Group Method with Arithmetic mean method. Taxa labels were arranged as follows: scientific name, genotype name and ploidy. Ploidy information was not mentioned for the species with undetermined ploidy. The scale bar represents the length of the tree branch.

### Development of Yam Microsatellite Markers Database

We built up a comprehensive database by integrating SSR markers and whole genome assemblies and annotations datasets (Figure 1). The Y2MD page contains data of four *Dioscorea* species and over 864,128 primer pairs. Also all SSRs data are downloadable in Excel, PDF, text, CSV format. The graphical user interface (GUI) menu has six tabs “ Home”, “ Species”, “ Tools”, “ SSRs”, “ Download “ and “ About “.

The “Species” tab displays information on each species with the possibility to choose the SSR of choice according to the type of patterns, the number of repeats, the position, etc. Information on experimentally validated SSRs and cross-species transferable SSRs are available in the “SSRs” tab. On the “Download” page, the links to the different genomes are available.

To enable the database user to perform some fundamental analyses, several functionalities were embedded in Y2MD among which JBrowse (**Figure 6a**), *insilico*PCR (**Figure 6b**), SSR Finder (**Figure 6c**), and Y2MD Blast (**Figure 6d**). Thus, Y2MD empowered the user to find an SSR with SSR Finder, and proceed to a quick amplification check with *insilico*PCR. Besides, the user can also blast his own sequence of interest onto a genome with Y2MD Blast tool. With JBrowse, the user is able to dive into the genome annotation map to find out potential genes related to a targeted SSR. Altogether, Y2MD offers a dynamic platform to the yam breeder to remotely use the available genomic resources in a GUI user-friendly manner. The website is freely accessible at http://yamdb.42web.io/.

**Figure 6.**
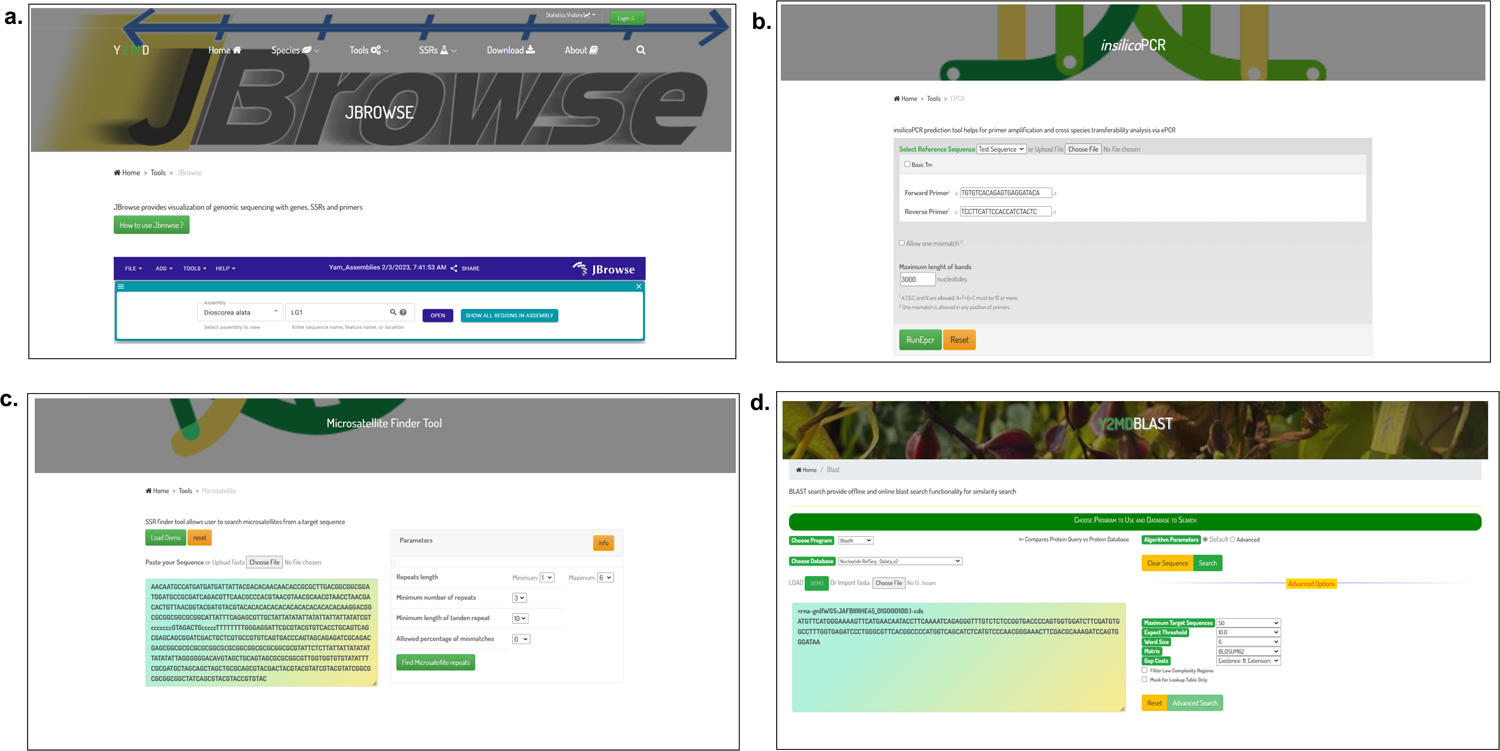
The yam microsatellite markers database: Y2MD. The website is freely available at http://yamdb.42web.io/. The web interface of Y2MD showcasing JBrowse (a), *insilico*PCR (b), SSR Finder (c), and Blast (d) functionalities.

## Discussion

### Developing genome-wide SSR markers in Dioscoreaceae

Microsatellite markers proved to be effective to characterise yam germplasms worldwide (Obidiegwu et al., 2009a; Obidiegwu et al., 2009b; Siqueira et al., 2011; Tostain et al., 2006; Tamiru et al, 2015; Arnau et al., 2017; Cao et al., 2021). The present study enabled the extraction of quite high numbers of microsatellites in different yam genomes. The extracted SSRs were more concentrated in the telomeres as reported by Jian et al. (2021). *Dioscorea zingiberensis* (253,856 microsatellites) had fewer microsatellites as compared to the other yam species. Nonetheless, we observed a high correlation (R^2^ =0.99) between the SSR numbers and chromosome lengths (**Figure 3)**. Our findings are similar to the report of Dossa et al. (2017) in sesame where the relationship between chromosome length and the number of SSRs on each chromosome showed a strong correlation (R^2^ = 0.94). Overall, the SSR densities observed in yam species (from 529 to 662 SSR/Mbp) are similar to that of sesame (507 SSR/Mbp) while, lower than other species such as Arabidopsis (*Arabidopsis thaliana*) and rice (*Oryza sativa*) (875 and 807 SSRs/Mbp, respectively) (Lawson and Zhang, 2006; Dossa et al., 2017). Comparing two genome assemblies of *D. alata* (Kabusa and TDa95/00328) highlighted major differences in terms of SSR numbers and densities. We speculate that the completeness of the genome assemblies (**Figure 2**) and the genetic distances between the two accessions may explain these differences. Among the observed motifs, mononucleotides that account for 50% of SSRs are the most represented (**Figure 3)**. However, they are not ideal targets for PCR markers (Clarke et al., 2001). The most abundant types of SSRs are mono (A/T)n, di (AT/TA)n, poly(ATA/ATT/TAT/TAA)n and therefore rich in AT. Similar results are obtained in potato (*Solanum tuberosum*) (Jian et al. 2021), Arabidopsis (Cavagnaro et al., 2010), sesame (Uncu et al., 2015), sorghum (*Sorghum bicolor*) (Yonemaru et al., 2009) and rice (Mccouch, 2002).

### Cross-species transferability of the SSR markers

The identification of polymorphic SSR markers will support research on yam genetics and breeding, especially in developing countries. At the genome scale, we identified 1170 cross-species transferable markers in the studied four Dioscorea species. Previously, Tamiru et al. (2015) showed that cross-amplification of SSR markers from *D. cayenensis*, revealed that 94.4% and 56.7% could be successfully transferred to *D. rotundata* and *D. alata*, respectively. Transferring markers between species is often considered a better investment than developing and testing new markers, especially when available funding is limited (Fan et al., 2013). Because in most breeding programs, different yam species are grown and evaluated together, therefore having SSR marker sets applicable on different species is highly useful. We also noticed that the number of cross-species transferable markers decreased significantly when *D. zingiberensis* was included (**Figure 4)**. Indeed, Bredeson et al. (2022) compared *D. alata* reference genome with the genomes of *D. rotundata* and *D*. *zingiberensis* and revealed substantial conservation of chromosome structures between *D. alata* and *D*. *rotundata*, but at a lesser extent with *D. zingiberensis*.

When several genome assemblies are available within the same species, *in silico* detection of polymorphic markers is resource-efficient in terms of time and cost. This approach has been successfully applied in sesame, spinach, and *Perca fluviatilis* (Dossa et al., 2017; Bhattarai et al., 2021; Xu et al., 2022). We compared the common SSR markers from the genomes of *D. alata* (TDa95/00328 and Kabusa) and obtained 93,92% polymorphic SSR markers. In other species such as potato, tomato and bell pepper, the levels of intraspecific polymorphisms appear to be rather limited (Stàgel et al., 2008), therefore much efforts were needed to develop informative SSR markers in these species. In *D. alata*, the high SSR polymorphic rates could be attributed to the dioecy nature of the species.

### Validating selected SSRs

In *Dioscoreaceae*, very few SSR markers have been experimentally (wet-lab) validated (Obidiegwu et al., 2009a; Obidiegwu et al., 2009b; Andris et al., 2010; Siqueira et al., 2011; Tostain et al., 2006; Tamiru et al., 2015; Arnau et al., 2017; Cao et al., 2021), most of which are from *D. alata* and *D. rotundata*. In other species such as rice (McCouch et al., 2002), sesame (Dossa et al., 2017), and cowpea (*Vigna unguiculata*) (Jasrotia et al., 2019), a large number of markers have been identified, experimentally characterised and available for genetic studies and breeding. In this study, we extracted 92 markers from the literature (wet-lab validated markers) and most of them were found among the markers created in this study. As the e-PCR parameters do allow only two mismatches and one insertion for the primers, this could be partly responsible for the absence of PCR products for the missing wet-lab validated markers. This analysis also provided the exact locations of these SSRs in the yam genomes, which will enable researchers to select markers from different chromosomes in order to have a good discriminatory power.

Out of 18 selected SSR markers, 17 were able to amplify by PCR several yam genomes in this study. The experimentally tested markers were from different chromosomes. They demonstrated a robust discriminatory power by clearly delineating the studied yam species and ploidy levels (**Figure 5**). In yam species, the basic chromosome number is 20 and most species harbour different cytotypes (Scarcelli et al., 2005; Arnau et al., 2009; Abraham et al., 2013; Sughihara et al., 2021). Prior to our work, *D. rotundata* cultivars were reported to be mainly diploid using flow cytometry, isoenzymes, DArTseq SNP and a set of six microsatellite markers (Gatarira et al., 2021; Scarcelli et al., 2005). Both *D. rotundata* cultivars investigated in this study were found to be diploid. Besides, the tested SSR markers also exhibited a promising accuracy even at intra-species level with a clear classification of diploid and tetraploid forms of *D. alata* cultivars. Based on the allele numbers, we report that yam species including *D. transversa*, *D. abyssinica*, *D. cayenensis*, and *D. nummularia* have cultivars with tetra- or higher ploidy levels (**Table 1**) (Gatarira et al., 2021; Sughihara et al., 2021). Therefore, we provided in the present study, SSR markers that can be used at low cost for ploidy check and species differentiation in yams.

### Deployment of Yam Microsatellite Markers Database for yam breeders

Many databases dealing with SSR marker data from various species have been made available to the public. However, no database is available to study yam microsatellites. A yam microsatellites marker database will be a useful platform for advancing genetic evaluation, genomic research, and yam breeding. The Pan Species Microsatellite Database (Du et al., 2020) containing SSR data from 18,408 organisms is limited by the lack of features such as ePCR, Jbrowse (Buels et al., 2016) and Blast (Altschul et al., 1990). This is also the case for the Tea Microsatellite Database (Dubey et al., 2020) and SSRome (Mokhtar and Atia, 2019). In this study, we compiled all generated microsatellite data into Yam Microsatellite Markers Database (Y2MD) (**Figure 6**). Y2MD is embedded with various useful tools such as JBrowse, Blast, SSR Finder and *insilico*PCR. These tools will facilitate the localization, SSR detection and polymorphism analysis for existing and new experimental yam SSR markers/nucleotide sequences. Y2MD has been developed to be a very user-friendly database enabling all users, especially those with limited knowledge or resources in bioinformatics to analyse their SSR related data.

## Conclusion

In this study, a graphical interface containing the SSR markers of four commercially important Dioscorea species was constructed and made available to the public via http://yamdb.42web.io/. To perform the database construction, microsatellites of four *Dioscorea* species were first extracted and primers were designed on the flanking sequences of these microsatellites. In total 1,201,570 microsatellites were extracted from all species with 864,128 designed primer pairs. Then, additional analyses were performed such as cross-species transferability and determination of polymorphic markers which are determining factors in plant genotyping. Experimental validation of selected markers were also performed to provide credence to our datasets. Overall, the SSRs developed in this work will be useful for genetic characterization of yam germplasms, quantitative trait loci (QTL) mapping and marker assisted breeding.

All of this information has been compiled into Y2MD and will be of great use for the yam breeding community, especially in developing countries. With the rapid development in genome sequencing projects, we expect more yam genomes to come. Hence, we will expand Y2MD to include new *Dioscorea* species as soon as their genome sequences are available. In addition, Y2MD will be enhanced by adding new tools and functionalities such as the integration of Primer3, introducing molecular markers associated with QTLs and new wet-lab validated SSRs.

## Availability of Data and Materials

All SSR data generated and the genomes used are available online at http://yamdb.42web.io/. The genome sequence of *Dioscorea alata* cultivar Kabusa has not been made available publicly yet, but it can be shared upon request to the corresponding author.

## Author Contributions

*Conceptualization*: Komivi Dossa, Diaga Diouf, Hâna Chair. *SSR validation experiment*: Ronan Rivallan, Erick Malédon, Hâna Chair, Komivi Dossa. *Data acquisition, curation and formal analysis*: Moussa Diouf, Yedomon Ange Bovys Zoclanclounon, Pape Adama Mboup, Diaga Diouf, Komivi Dossa. *Database building*: Moussa Diouf, Pape Adama Mboup, Komivi Dossa. *Funding acquisition*: Komivi Dossa. *Writing ± original draft*: Moussa Diouf, Yedomon Ange Bovys Zoclanclounon, Komivi Dossa. *Writing ± review & editing*: Diaga Diouf, Hâna Chair. All authors have read and approved the final version of this manuscript.

## Acknowledgment

Not applicable.

