## Supplementary material for "Genome-wide development of intra- and inter-specific transferable SSR markers and construction of a dynamic web resource for yam molecular breeding: Y2MD": Supplemtary_Figure.pdf

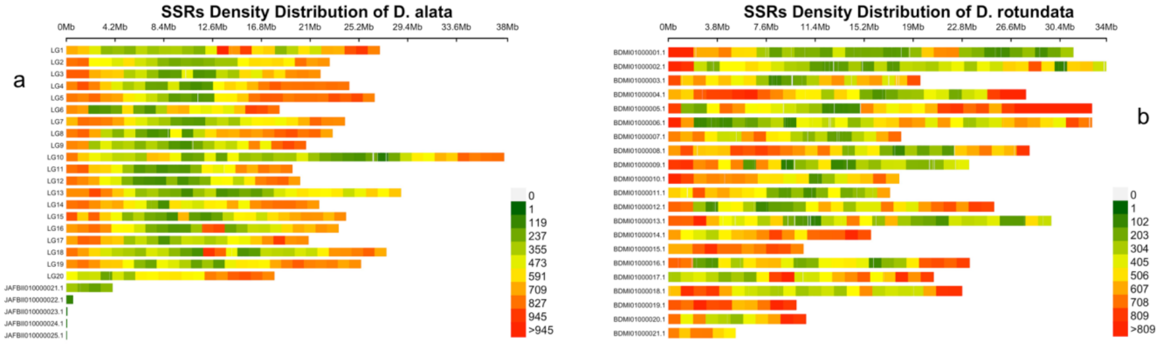


**Supplementary Figure 1**: Overview of the high-density SSR physical map in *Dioscorea alata (*a) and *Dioscorea rotundata (b)*. The bar represents the number of SSR markers within a 1-Mb window.
